# Meta-analyses reveal no clear demographic consequences of phenological change across taxa

**DOI:** 10.1101/2025.10.29.685396

**Authors:** Elsa Godtfredsen, Paul CaraDonna, Alicia Foxx, Justin Bain, Brendan Connolly, Kathryn Dawdy, Alissa Doucet, Jacquelyn Fitzgerald, Gwen Kirschke, Jane Ogilvie, Ceci Rigby, Joshua Scholl, Dan Sandacz, Samantha Rosa, Alexandra Zink, Amy Iler

## Abstract

The timing of life-cycle events (phenology) is key to organism ecology and success. Climate change is shifting phenology to earlier dates globally, but generalizable trends of how phenological change impacts demography are still unknown. Therefore, we conducted a meta-analysis to quantify the effects of interannual phenological variation and phenological shifts on demographic vital rates (survival, growth, and reproduction). Our dataset includes 138 taxa from 83 studies, representing different study approaches (observational and experimental) for plants and animals. Using these data, we asked three primary questions: 1). How does phenological variation affect demographic vital rates? 2). Are directional shifts in phenology predictive of changes in demographic vital rates? 3.) Do relationships between phenology and demography depend on taxa, vital rate, and study type? For studies of phenological variation, earlier events conferred demographic benefits whereas later events were associated with demographic costs, with most of the evidence coming from bird and reproduction-focused studies. In contrast, directional phenological shifts were not predictive of demographic responses over time or in experiments. While there was evidence that phenological events shifted earlier through time, there was not significant change in demographic vital rates over those same time periods. These results are consistent with the hypothesis that organisms may be able to track environmental conditions to maintain demographic performance by shifting their phenology to earlier dates under climate change. Critically, our meta-analysis clarifies that while earlier phenological events tend to confer demographic benefits in the context of phenological variation, directional phenological shifts to earlier timing did not show demographic benefits.

## INTRODUCTION

Investigating broad demographic effects of organismal timing (phenology) is vital because phenological shifts are among the most widespread signals of climate change^1–7^. Phenology is at the center of an organism’s ecology and evolution and dictates the abiotic and biotic conditions to which organisms are exposed, thereby affecting survival, growth, and reproduction^8–13^. While phenology varies from year to year due to interannual variation in environmental cues, climate change has triggered shifts to earlier life cycle events in a variety of taxa^14–22^. It is common in the literature to use performance in earlier years to predict future climate change performance, but interannual variation and phenological shifts will likely have different demographic impacts. Despite widespread hypotheses that phenological change has strong demographic consequences, generalizable conclusions are still limited, and we do not know whether positive, negative, or neutral effects are the norm^23^. Therefore, we conducted a meta-analysis to quantify the effect of variation in phenology and phenological shifts on demographic consequences.

The demographic consequences of phenology can depend on the ability of an organism to phenologically “track” its environment; ideal tracking would allow an organism to experience its original conditions in response to an environmental change^10,24^. We hypothesize that tracking under climate change would result in a phenological shift *without* strong effects on demographic vital rates; this is because the change in timing allows the organism to experience similar environmental conditions (on a different calendar date). However, tracking one environmental cue does not always account for changes in other factors, which can result in exposure to novel conditions^8^. Phenological shifts can alter an organism’s growing season length^25,26^, temperature exposure^27,28^, and access to resources^29,30^, all of which can confer demographic benefits or costs, depending on the organism and the environment.

Any reported demographic consequence of phenology will likely depend on several factors, including taxa studied, demographic vital rate(s) measured, or study approach (e.g., experimental or observational). Taxonomic groups can differ in their magnitude of phenological change^6,31,32^. For example, plants display larger phenological change than other organisms in some ecosystems and may experience more intense demographic consequences compared to mobile organisms^6,33^. Which demographic vital rate is measured can also affect the magnitude of the demographic consequence and past synthesis suggests that phenology has stronger impacts on reproductive vital rates than growth and survival^34^. Lastly, demographic outcomes in experiments might differ from observational time series because experimental treatments can introduce artifacts (such as drought due to warming treatments) that can impact demography independently of changes to phenology^35^.

To explore the demographic consequences of phenological change we ask the following questions: 1). How does interannual phenological variation affect demographic vital rates? 2). Are directional shifts in phenology predictive of changes in demographic vital rates? 3.) Do relationships between phenology and demography depend on taxa, vital rate, and study type? Without a quantitative understanding of how phenological change affects demographic vital rates, we are missing key knowledge of how climate change affects biodiversity.

## METHODS

We used a meta-analysis to investigate potential demographic consequences of phenological variation and shifts in phenology. Our final dataset included 138 species from 83 studies. The dataset contained mostly terrestrial studies from the northern hemisphere; most studies were conducted on birds (38%) and plants (38%) compared to other taxa (Table S1).

### Literature sources & effect size estimates

We sourced studies from the narrative review conducted by Iler et al. (2021) and an updated search conducted in May of 2024 using the same search terms and protocols. In brief, papers needed to have the ability to document a phenological shift through time or in response to climate and to have measured a demographic outcome (e.g., reproduction, survival, etc.) (see details in Appendix 1). Out of the 406 papers gathered from the initial and updated searches, 83 studies met the criteria to be included in the final meta-analysis.

We extracted effect estimates from each study according to the three distinct data types illustrated in Figure 1: (i) phenological variation, (ii) shifts across time, and (iii) shifts across treatments (Figure 1). (i) Most phenological variation estimates (76%) consisted of a single slope that captured a change in a demographic vital rate in response to phenological variation across multiple years (Figure 1). Wilson & Martin (2010) assessed the responses of two bird species to spring weather variation and regressed the breeding date against reproductive output over four seasons. A limited amount (14%) of phenological variation estimates consisted of a slope that captured a change in demographic vital rate in response to phenological variation across experimental treatments. (ii) Shifts across time estimates consisted of two slopes that represented independent changes in both phenological and demographic metrics through time (Figure 1). Macgregor et al (2019) separately assessed the date of emergence and population abundance over the same 20-year period in 130 bird species. (iii) Shifts across treatment estimates included raw means of both phenological and demographic metrics in response to experimental treatments (Figure 1). Pérez-Ramos et al (2020) applied warming and drought treatments to multiple plant species in a Mediterranean ecosystem and assessed how treatments affected peak flowering and flower number.

**Figure 1.**
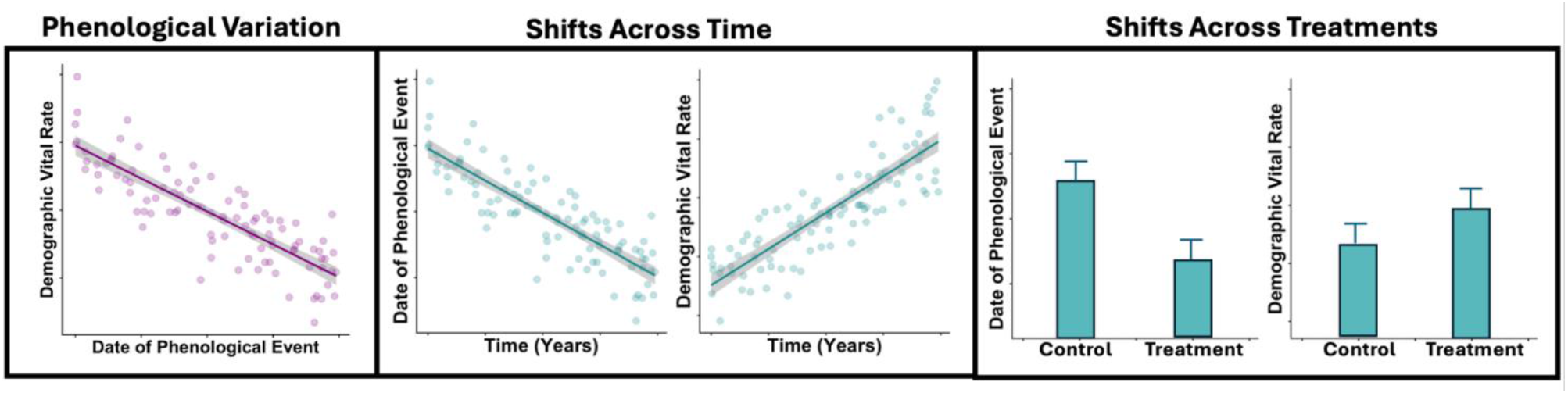
Types of Source Data Extracted For Analyses. Phenological Variation. For these studies we extracted both the slope of the regression line (Date of Phenological Event X Demographic vital rate) and its standard error (SE). Sources of phenological variation included multiple years as well as experimental treatments. **Shifts Across Time**. For these studies, we extracted both the phenological slope (Time X Phenology) as well as the demographic slope (Time X Demography) and SE’s for both relationships. **Shifts Across Treatments**. For these studies, we extracted both the mean and SE of the date of the phenological event as well as the demographic vital rate in control and treatment.

We also extracted an associated measure of variation for every estimate (standard error (SE), standard deviation (SD) or confidence intervals (CI), depending on what was reported). We then standardized the measures of variation by converting them into SE. Additionally, we altered treatment categorization of the experimental estimates to be consistent with climate change scenarios (seven studies altered). For example, if an experiment manipulated snowpack, we categorized the reduced snowpack treatment as the treatment, and the unmanipulated treatment as the control. In contrast, if an experiment added snow, then we categorized the added snow as the control and the unmanipulated treatment as the treatment. This ensured that responses to experimental manipulation were indicative of climate change. We also recorded the statistical model type and error families used to calculate each estimate. Additional parameters for paper inclusion, details of data extraction, and common reasons for exclusion can be found in Supplemental Materials (Appendix 1, Table S2).

### Statistical Analyses

Our goal was to assess how phenological variation affected demographic vital rates and to investigate if phenological shifts predicted demographic change (across time and treatments). All analyses were conducted in R (V4.4.2)^36^.

#### Phenological Variation

The phenological variation dataset spanned multiple taxa, including birds (46%), plants (33%), fish (10%), mammals (8%) and amphibians (1%). Most estimates (*i*.*e*., slopes) focused on reproduction as the demographic response (79%), but some estimates also focused on size and growth rate (9%), survival (8%) or abundance (4%). The average study length was 15.04 ± 13.10 years.

We grouped the slopes and SE’s based on the error family used to calculate them in the original study (Gaussian, binomial, Poisson) and analyzed them separately because their effect sizes are indicative of responses on different scales. We used random effects models (*rma*.*mv* function)^37^, which assume that the true effect size is different across studies and weights estimates by their inverse variance; we included study ID as a random effect to account for non-independence among estimates^38^. Lastly, we used robust error estimation (*robust* function)^37^ to calculate our meta-estimates and CIs because it further controls for non-independence between multiple estimates from a singular study^39^.

We constructed ‘full’ models for each error family grouping (Gaussian, binomial, Poisson) that included all studies in each category. To understand if there were different trends in specific taxa or demographic vital rates, we conducted separate analyses when sample size was adequate (over 5 studies and 10 effect sizes per group). Due to limited sample sizes in many groups, we analyzed studies that focused on reproduction, survival and size, plants, animals, and birds separately. We used migratory status as a modulator (meta-analytic interactive term) in our bird analyses to investigate evidence for differences in phenological responses in migratory vs. non-migratory birds^40^.

#### Shifts Across Time & Treatments

Time series studies were often on birds (53%), followed by insects (22%), mammals (16%), fish (5%), plants (2%), and amphibians (2%). Most studies focused on reproduction as the demographic response (71%), but some studies also focused on abundance (22%), or size and growth rate (7%). The average study length was 30.53 ± 14.56 years. To analyze the trends for both phenology and demography over time, we used two random effects models (*rma*.*mv* function)^37^ to assess if there was a generalizable trend for phenology (earlier or later) and demography (cost or benefit). Next, we used linear mixed effects models (LMM, *lmer* function)^41^ to assess if changes in phenology over time were predictive of changes in demography over the same time period. The LMM consisted of the slope of phenological change across time as the *predictor* and the slope representing demographic change over the same time period as the *response*, with study ID as a random intercept term. Including study ID as a random intercept term accounts for most studies having multiple estimates, either due to the inclusion of multiple species or multiple demographic vital rates.

We constructed ‘full’ models that included all the shifts across time studies. To understand if there were different trends in specific taxa or demographic vital rates across time, we conducted separate analyses when sample size was adequate. Due to limited sample sizes in many groups, we analyzed studies that focused on reproduction and birds separately. To determine if migration status modulated the effect of phenological shifts, we included migration status (yes or no) as an interactive term for bird-specific analyses. To assess whether results depended on whether significant phenological shifts had occurred, we repeated LMM regressions only for cases in which the phenology estimates 95% CI did not cross 0 (phenology was significantly advancing or becoming later over the time period)^42^.

In contrast to the shifts across time dataset, shifts across treatment studies were mostly on plants (91%), with low representation (∼1-2% each) of reptiles, birds, insects, amphibians and mammals. Most studies focused on reproduction as the demographic response (79%), but some studies also focused on size and growth rate (12%), or survival (9%). The average study length was 1.85 ± .76 years. To analyze the shifts across treatment studies, we first calculated effect sizes that represent the change in both the phenological variable and demographic variable in response to the treatment using the log response ratio (lnRR)^43,44^. We calculated two lnRRs for each reference mean and treatment mean pair for phenology and demography using the following equation^43^:

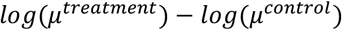

We calculated the standard error for each lnRR using the following equation:

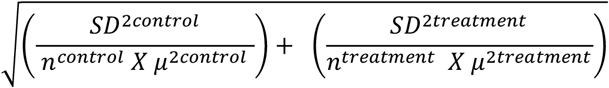

All but two experimental studies had multiple estimates per study due to assessment of multiple species, different demographic responses, or multiple treatments. To account for this, we used LMMs with the lnRR of the phenological metric as the predictor and lnRR of the demographic metric as the response and study ID as a random intercept. We first constructed a ‘full’ model that included all shifts across treatment studies and then conducted separate analyses on plants, reproduction, survival and size, Treatment-Warming, Treatment-Snow and Treatment-Other. To understand if plant traits modulated the relationship, we included growth form (woody or herbaceous) and life history strategy (annual or perennial) as interactive terms for plant-specific analyses. Additionally, we repeated LMM regressions only for cases in which the phenology lnRR 95% CI did not cross 0 (phenology was significantly advanced or later in the treatment relative to the control)^42^.

#### Publication bias assessment

To check for potential bias within our datasets, we utilized contour enhanced funnel plots (*funnel* function)^37^, which allow one to visualize possible asymmetry across gradients of significance^45^ and Egger’s regression to test for funnel plot asymmetry (*regtest* function)^37,46^.

## RESULTS

Our meta-analysis of 83 studies, representing 128 taxa, including plants and animals, revealed a fundamental distinction in the demographic consequences of phenological change. In studies of phenological variation, earlier events were associated with demographic benefits. In contrast, directional phenological shifts toward earlier events were not predictive of directional demographic consequences.

### Phenological Variation

Overall, there was a significant negative relationship between phenology and demographic vital rates, implying that earlier events conferred demographic benefits while later events conferred demographic costs (Table 1; Fig 2). This general negative trend was consistent across taxa and demographic vital rates but was only statistically significant for the full model, reproduction, and birds (Table 1; Fig. 2). Migration status did not significantly modulate the relationship between phenological variation and demographic consequences in birds (Table 1). Plant studies exhibited the largest meta-estimate but also the greatest amount of variation (Table 1; Fig. 2). Across all model types, (Gaussian, binomial and Poisson), results were consistent (Table 1, Appendix 2, Figure S1, Figure S2).

**Table 1.**
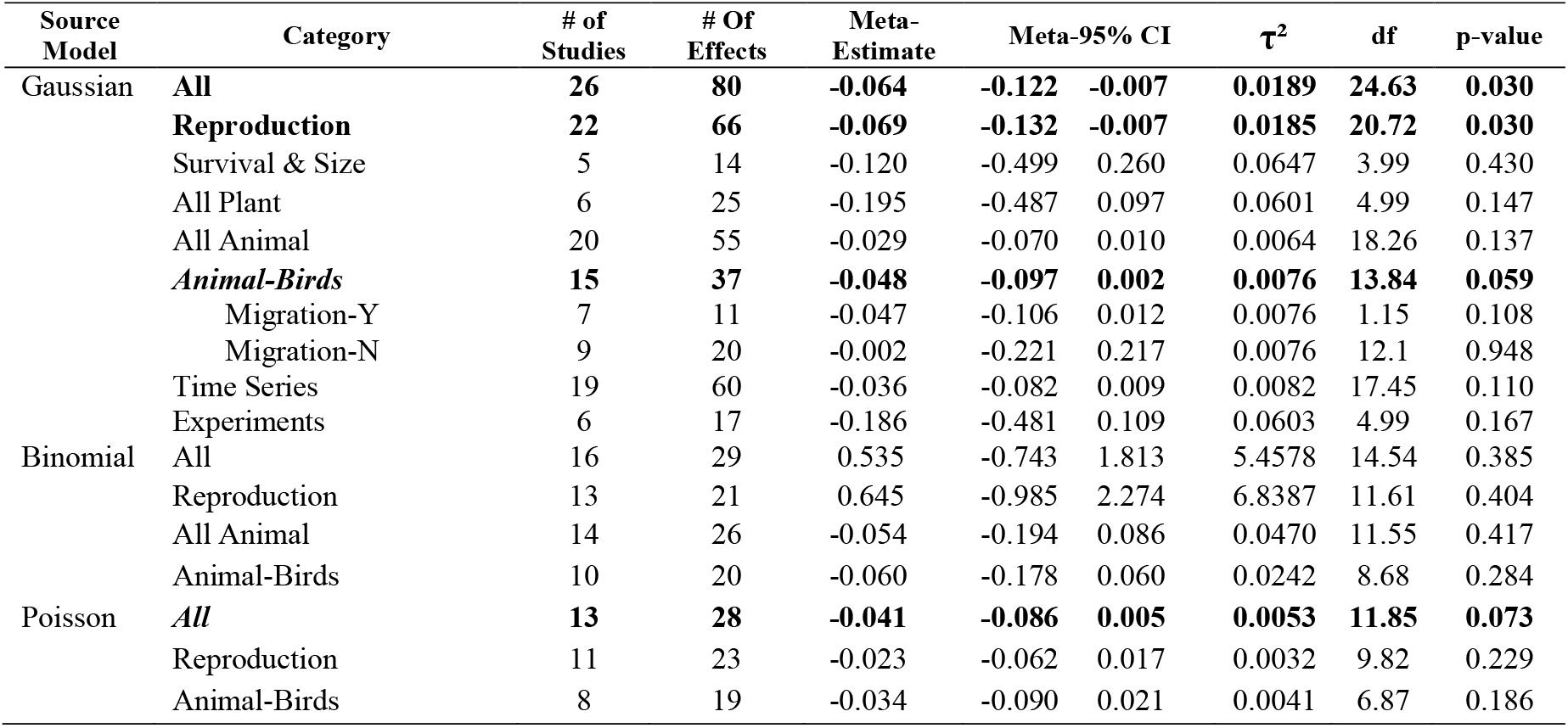
Results of phenological variation studies meta-analyses. All results came from rma.mv models from the metafor package with study ID as a random effect. Sample sizes (both the number of studies and the number of effect sizes included to account for studies with multiple effect sizes), the meta-estimate and meta-CI (calculated with robust error estimation) are shown for each analysis. Presented analyses all include more than five studies and 10 effect sizes (if groups were lower than these sample sizes, these were excluded from separated analyses but are included in the “All” category). Analyses that show marginally significant (p-value ≤ 0.1 and ≥ 0.05) or significant (p-value ≤ 0.05) are bolded. τ^2^ represents the between study variation in effect sizes, a 0 τ^2^ value indicates that no true variation in effect sizes was detected and all variation is the result of estimation error. Estimated df is also presented.

**Figure 2.**
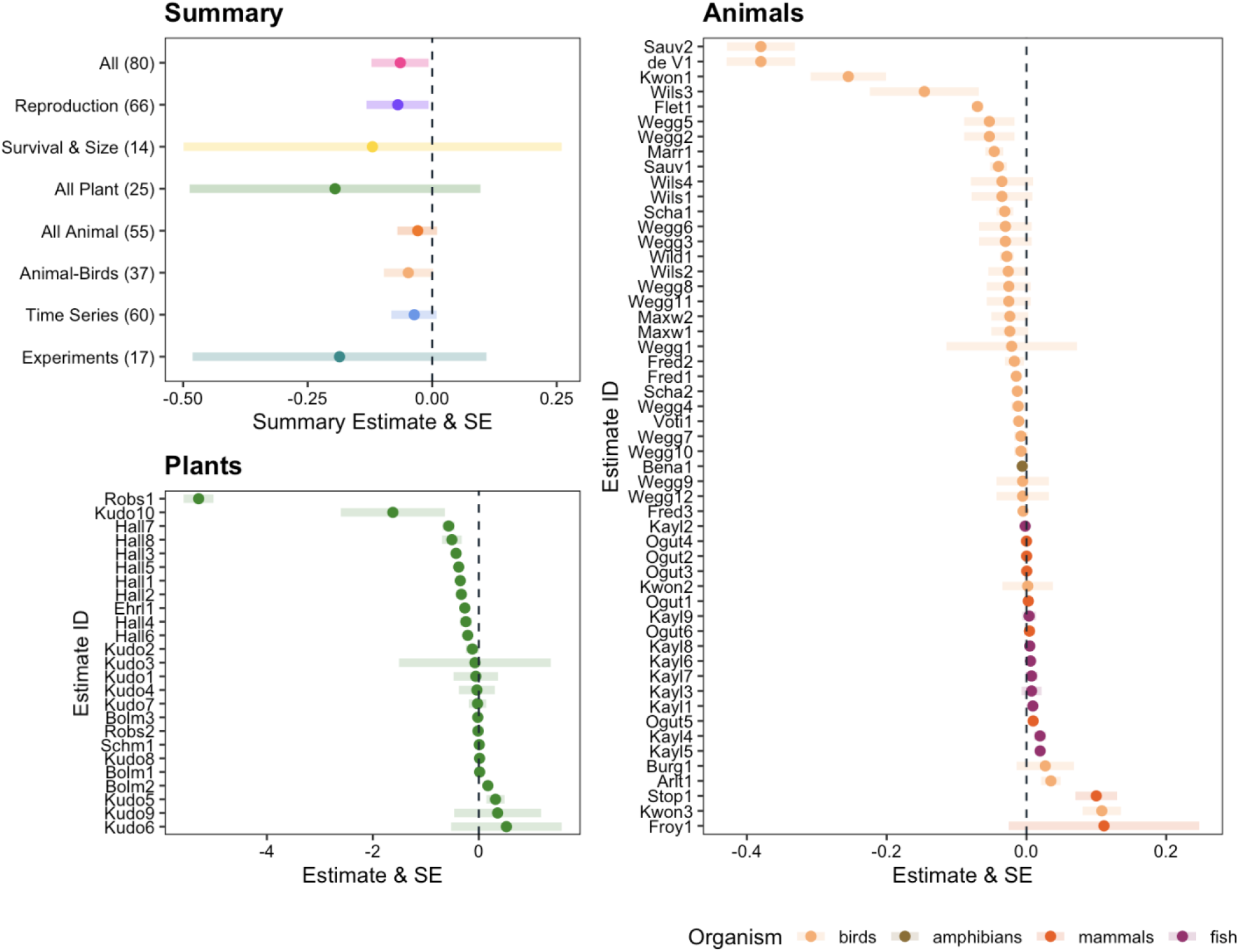
Meta-Estimates & Raw Estimates of Phenological Variation Studies. Plant and animal estimates are visualized on separate panels due to large differences in scale of estimates. The meta-estimates and meta-CI’s calculated for each group of studies are shown with color corresponding to group. The number of effect sizes included in the analyses are presented in parentheses, the central dot represents the meta-estimate for this group and the bars represent the lower and upper summary meta-CI for the group, with a dotted line indicating 0. **Plants:** The raw estimate and SE for each estimate (multiple estimates for many studies) are shown. Study ID is shown on the y axis (including first four letters of first author name and estimate number within the study). All estimates are visualized with error bars indicating SE although some are not visible due to large scale differences between studies. **Animals:** The raw estimate and SE for each estimate (multiple estimates for many studies) are shown. Study ID is shown on the y axis (including first four letters of first author name and estimate number within the study). All estimates are visualized with error bars indicating SE although some are too small to be visible due to large scale differences between studies. Colors indicate organism type.

**Figure 3.**
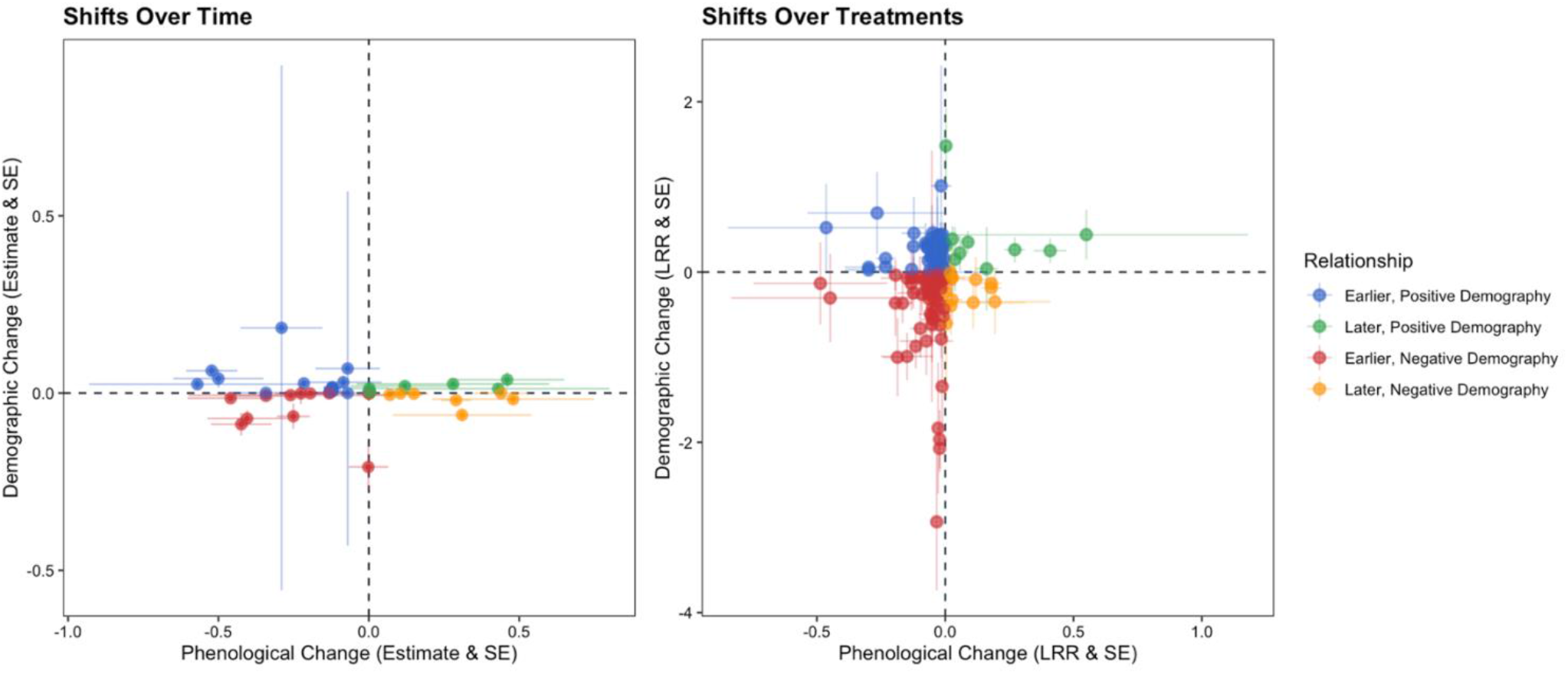
Slopes of Shifts Across Time & lnRR of Shifts Across Treatments. Y-axis differentiation maintained for clarity of data; note scale difference on Y-axis between panels. **Shifts Across Time** studies are presented by plotting estimates and SE for change in phenology across a time period plotted against estimates and SE for change across demography in the same time period. **Shifts Across Treatments** studies are presented by plotting the lnRR (log response ratio that presents the ratio of change from treatment to control) of the phenological metric against the lnRR of the demographic metric. Each dot represents one of these relationships and the error bars represent the SE for each lnRR. Colors coordinate with quadrants that represent relationships between the slopes and lnRRs.

We assessed these datasets for publication bias using Egger’s regression (Table S5). There was significant asymmetry detected for the full, reproduction, survival and size and experimental datasets (Table S5). While this can indicate publication bias, asymmetry is also a common artifact of high heterogeneity. We found that all asymmetric datasets combined plants and animals, two groups with large differences in their reported effect sizes. Notably, when plants and animals were analyzed separately, no significant asymmetry was detected (Table S5). This strongly suggests the observed asymmetry is driven by high (and valid) heterogeneity between taxa rather than by publication bias.

### Shifts Across Time

Phenology tended to shift to earlier dates over time, but this was generally not predictive of demographic consequences over the same period. There was a marginally significant trend for earlier phenology over time (meta-estimate = *-0*.*1245, meta-95% CI= -0*.*260 – 0*.*011, t= 0*.*042, p= 0*.*069*), and no significant trend for demography over the same period (meta-*estimate* = 0.0005, *meta-95% CI = -0*.*004 – 0*.*005, t = 0*.*001, p-value = 0*.*826*). All relationships between phenological shifts and demographic responses are negative (earlier phenology associates with demographic benefits) and nonsignificant in our LMMs (Table 2; Fig. 2). The significance of these relationships is consistent when we restrict the analyses to include only studies that reported significant phenological shifts (95% CI does not cross 0) (Table S4). There was no asymmetry detected in the time series datasets (Table S5).

**Table 2.** Results of LMMs to understand the effect of phenological shifts on demographic vital rates across time and treatments. All results came from LMM models constructed in lme4 with study ID as a random effect. Sample sizes (both the number of studies and the number of effect sizes included to account for studies with multiple effect sizes). Presented analyses all include more than five studies and 10 effect sizes (if groups were lower than these sample sizes, these were excluded from separated analyses but are included in the “All*”* category). Analyses that show marginally significant (p-value ≤ 0.1 and ≥ 0.05) or significant (p-value ≤ 0.05) are bolded. Historical comparisons and comparisons of extreme years are excluded from all experimental analyses except for Treatment-other.

| Data Source | Category | # of Studies | # Of Effects | Estimate | 95% CI |  | *t* | p-value |
| --- | --- | --- | --- | --- | --- | --- | --- | --- |
| Shifts Across Time | All | 13 | 45 | -0.025 | -0.090 | 0.043 | -0.73 | 0.383 |
|  | Animal-Birds | 7 | 24 | -0.030 | -0.161 | 0.101 | -0.45 | 0.652 |
|  | Migration Status | 7 | 24 | -0.106 | -0.382 | 0.169 | -0.72 | 0.472 |
|  | Reproduction | 10 | 38 | -0.053 | -0.171 | 0.079 | -0.87 | 0.386 |
| Shifts Across Treatments | All | 26 | 132 | 0.194 | -0.495 | 0.881 | 0.55 | 0.580 |
|  | Plant | 22 | 126 | 0.202 | -0.508 | 0.911 | 0.56 | 0.576 |
|  | Growth Form | 22 | 126 | 4.088 | -0.950 | 9.130 | 1.58 | 0.185 |
|  | Life History | 22 | 126 | 0.434 | -1.114 | 1.999 | 0.55 | 0.586 |
|  | Reproduction | 22 | 104 | 0.109 | -0.647 | 0.862 | 0.28 | 0.776 |
|  | **Survival & Size** | **8** | **28** | **4.133** | **1.859** | **6.655** | **3.55** | **>0.001** |
|  | Treatment-Warming | 15 | 70 | -0.411 | -1.266 | 0.449 | -0.96 | 0.344 |
|  | Treatment-Snow | 8 | 41 | 0.821 | -0.985 | 2.631 | 0.88 | 0.382 |
|  | Treatment-Other | 11 | 49 | -0.154 | -0.657 | 0.373 | -0.59 | 0.552 |

### Shifts Across Treatments

Consistent with shifts across time, experimental treatments generally prompted a shift to earlier phenology, but this did not incur consistent demographic responses. We observed a significant trend for earlier phenology in treatments in comparison to controls (meta-estimate= *-0*.*0435, meta-95% CI= -0*.*071 – 0*.*016, p= 0*.*003*), and no significant trend for demography in the same treatment and control comparisons (meta-estimate= - *0*.*0801, meta-95% CI= -0*.*187 – 0*.*026, p-value: 0*.*134*). The direction of the relationship between phenological and demographic change in experiments was inconsistent (Table 2, Fig 2). While survival and size showed a significant, positive relationship, all of the included studies had earlier phenologies in the treatment relative to the control (Table 2, Figure 2). In this context, a positive relationship indicates that in treatments with vastly earlier phenology than controls, there was a negative impact on survival and size, while treatments that were slightly earlier than controls had both positive and negative impacts on survival and size.

The significance of the relationships is consistent when we restrict the analyses to include only studies that reported significant phenological shifts (95% CI does not cross 0) (Table S3). There was significant asymmetry, which can suggest publication bias, detected in the experimental demography dataset but not in the experimental phenology dataset (Table S5). Experimental datasets may be more prone to publication bias than observational studies due to the challenge of mimicking climate change using experimental treatments. For example, if experimental treatments do not result in strong effects on demography, those studies may be less likely to be published than observational studies, due to the assumption that the treatments were not ‘extreme’ enough to mimic climate change scenarios.

## DISCUSSION

Our meta-analyses suggest that phenological shifts are not predictive of demographic consequences across time nor treatments, but that there are demographic benefits to earlier timing in the context of phenological variation. Our study clarifies that demographic consequences of phenology can differ depending on the type of phenological change studied (variation vs. directional shifts). Although demographic benefits might be detected in earlier years or treatments, this does not guarantee that demographic benefits will be sustained if there is a consistent population-level shift to earlier timing.

Our finding that earlier life-cycle events confer demographic benefits in the context of phenological variation scales up a well-documented individual-level process. Selection gradient studies assess the effect of individual-level phenological variation on demography and find that, within a given year, earlier individuals out-perform later individuals^47,48^. To our knowledge this is the first meta-analysis,, to demonstrate that this pattern holds true at the population level. While this population-level effect may seem intuitive, given that populations are composed of individuals, but documenting population-level demographic impacts is a necessary step. It provides the empirical basis for understanding how a shift in the distribution of phenology (i.e., “phenological variation”) translates to demographic consequences for the entire population.

The significant overall relationship in our phenological variation analysis was primarily driven by studies on birds (which were marginally significant, p=0.059) and reproduction. There are several possibilities for why there is a relationship between phenological variation and demography in birds in particular, including migration constraints^49^, harsh breeding conditions^50,51^, and reliance on insect prey with large abundance pulses^52–54^. We hypothesized that migratory status would be a key modulator, as migrants often show less phenological plasticity and their population declines have been linked to this inflexibility^40,55^. However, migratory status was not a significant modulator in our analysis; both migratory and non-migratory birds showed a similar trend of demographic benefits from earlier events. This suggests that while migrants may be less *plastic*, they face the same fundamental *selective pressure*: all birds experience demographic benefits when their phenology aligns with earlier conditions and costs when they miss this window.

In contrast, plants had the largest meta-estimate of any taxa but with the highest variability, which led to a non-significant relationship between phenological variation and demography. This high variability is likely attributable to two factors. First, their sessile nature may make them more sensitive to interannual variation in their immediate environment. Second, our dataset had a high proportion of plant studies that reported “flower number” as the demographic response, which is a highly variable measure of reproductive *effort* rather than true reproductive *output*. Therefore, this non-significant result should not be interpreted as a lack of an effect. Rather, the large magnitude of the variation itself indicates that plant populations are experiencing strong, though highly context-dependent, demographic responses to phenological variation.

Reproductive vital rates were the primary driver of the significant “phenological variation” trend ; survival and size vital rates, by contrast, showed no significant relationship. This is likely a direct consequence of a bias in the available literature. The majority of studies (88%) in our dataset linked reproductive *timing* (e.g., egg-laying date) with reproductive *outcomes* (e.g., number of offspring).Therefore, it is not surprising that reproductive vital rates,, that these studies found a significant relationship. While many studies reported multiple demographic vital rates, few reported the full suite of vital rates (reproduction, survival and size) which would allow us to assess potential demographic trade-offs^60–62^. We are missing the full picture of organism response and potential for effects on population dynamics without further demographic information. Therefore, we encourage further study of non-reproductive vital rates in response to phenological variation.

In contrast to the phenological variation results, there was not consistent evidence that phenological shifts were predictive of demographic consequences over time. Most studies showed populations shifting to earlier phenology, but these shifts were not predictive of demographic trends. This key finding is consistent with two primary hypotheses. The first is phenological “tracking**”** : organisms that successfully shift their timing to track their original environmental conditions should, by definition, maintain their demographic performance, resulting in no net demographic change. The second hypothesis is demographic decoupling, where phenology is not the only factor affecting demography over time. For example, warmer temperatures—the common driver of earlier events—can have multiple, conflicting consequences. A study on red grouse found that while warmer springs led to earlier breeding and larger clutches, these benefits were completely offset over time by an increase in tick-borne illnesses, which are also favored by warmer temperatures. Thus, the lack of a predictive relationship in our “shifts over time” analysis suggests that organisms are either successfully tracking their environment or that other climate change-driven factors are decoupling phenological shifts from their demographic outcomes.

Consistent with the shifts over time, experimental phenological shifts were also not predictive of demographic consequences. Here, the lack of a trend is likely due to the high proportion of plant-focused experiments (91%). As our phenological variation results showed, plants have highly variable demographic responses, with both positive and negative outcomes to earlier phenology. This inherent variability likely averages out any consistent directional trend in a broad meta-analysis.

It is also possible that experimental treatments had unintended side effects that could obscure the relationship between phenology and demography. For example, if a warming experiment induced both earlier flowering and drought stress^63^, the drought stress may have a larger effect on demographic vital rates than earlier flowering. That being said, a past meta-analysis found evidence for a link between phenological sensitivity to warmer temperatures and demographic benefits, specifically in plant warming experiments^10^. The seeming contrast in the findings of this meta-analysis with our own is likely due to differences in data; our dataset included non-warming treatments and focused on the direct relationship between phenology and demography rather than quantifying phenological sensitivity to temperature.

The high variability in plants is important to consider as it may obfuscate generalizable predictions for plant communities as changes in phenology become increasingly common. Therefore, we encourage more combined long-term monitoring of phenology and demography in plant communities, ideally combined with experimental manipulations that attempt to isolate the effects of phenology on demography.

The discrepancy between our variation and shift analyses strongly suggests that short-term demographic benefits from earlier phenology are not sustained over long-term directional shifts. This is not simply an artifact of using different datasets for each analysis; the pattern persists for birds, a group well-represented in both datasets. A plausible biological explanation for the discrepancy is costs of reproduction^60,66^. An organism might have ample resources to increase reproductive output in a single, isolated “early” year, continued higher rates of reproduction would come at the cost of growth or survival if this opportunity happens more consistently across years, as is the case with a directional phenological shift. Therefore, we should be cautious of using high rates of reproduction in earlier years as evidence of climate resilience, as such benefits might not be sustained over directional shifts in phenology.

Our findings are necessarily limited by the available literature, which is heavily biased toward temperate, terrestrial ecosystems and focused on plants and birds. To understand the global implications of phenological change, future work must expand to critically under-represented groups—such as amphibians, reptiles, and insects—and to aquatic and tropical habitats^34^. We advocate for more work that assesses the consequences of phenological variation and shifts on non-reproductive demographic vital rates such as growth and survival, especially for long-lived organisms in which these are the vital rates most likely to have population-level consequences^23,67–69^.

In conclusion, our results encourage caution in using demographic responses in early years as proxies for demographic responses of populations exhibiting phenological shifts, as is often done in the literature. Although it is clear that phenological shifts can have demographic consequences^8,23^, we found no generalizable trend in the direction of these responses. This suggests that as organisms shift their life histories in response to climate change, they are just as likely to experience demographic costs as they are to find benefits.

## Supporting information

Appendix 1, Appendix 2 and Supplementary Visuals

