## Appendix 1, Appendix 2 and Supplementary Visuals for "Meta-analyses reveal no clear demographic consequences of phenological change across taxa"

### Appendix 1: Supplemental Methods

#### Study Inclusion Parameters and Data Extraction Details

We sourced the papers for the meta-analyses from the review conducted by Iler et al in 2021 (Iler et al. 2021). The papers for this review were selected from search criteria from Web of Science applied to titles, author keywords and abstracts of articles: (∗synchron∗ OR phenolog∗ OR "temporal overlap" OR mismatch) AND "climate change" AND (demograph∗ OR reproducti∗ OR survival OR "vital rate∗") and papers were included that were published by August 2020. These search parameters returned 1,273 papers that were then assessed using two criteria: 1. Could the study quantify a phenological shift through time or phenological sensitivity to climate change? 2. Did the study measure a demographic consequence, including changes in vital rate (survival, growth, reproduction or recruitment), fitness, population abundance (number of individuals) or population growth rates (either per capita or finite increases in lambda). Only studies that incorporated field components were included, with purely theoretical and lab studies excluded. A total of 238 publications met these criteria and were included in the review by Iler et al. (2021).

In this study we only included papers where metrics could be extracted for each species separately. If papers only presented combined metrics for multiple species and authors were unable to be contacted, these papers were excluded. If a study included metrics for multiple populations or years separately (20% of all studies), we collected data on each population and year separately and controlled for pseudo-replication by using study ID as a random effect. We only included papers that used a frequentist framework for statistical analyses to ensure statistical consistency when analyzing effect sizes. If the error family used in models was unclear, authors were contacted for clarification. In the two cases where author contact was unsuccessful, we assumed a Gaussian distribution was used due to the compatibility of data type with this distribution.

When experimental studies included combined manipulations, such as heat and precipitation, we only collected data on the control and separated treatments and excluded the combined treatments. For papers that compared extreme years or historical periods, the warmer period was always specified as the treatment and the colder period as the control. We applied the same logic to snowmelt manipulations, where the earlier snowmelt situation was specified as the treatment and the later snowmelt as control to mimic expected climate change scenarios. Since many different measurements of variation are commonly used (standard deviation, standard error and confidence intervals), all measurements of variation were converted to SD and then used to calculate SE and variance to standardize measurements of variation for analyses.

**Protocols for Non-Reported Metrics**

When papers did not report the necessary metrics in text form, we followed a simple protocol. If the information (slopes, means or measures of variation) were represented in figures but not reported in text, we used the package meta-digitize in R (V4.2.2) to obtain necessary metrics. If metrics were unavailable in text or visual form, we contacted the authors to attempt to obtain the data. If the contact was unsuccessful, the paper was excluded from the meta-analyses. A total of 77 authors were contacted, with a 42% response rate.

### Appendix 2: Supplemental Results

*Binomial & Poisson Phenological Variation Results*

Studies using a binomial error distribution are dominated by birds (68%) and focus on survival and reproductive success. None of these relationships were significant (Table 1; Fig. S1). Studies using a Poisson error distribution are also dominated by birds (67%) and focused on reproductive counts such as clutch size and number of fledglings. The Poisson results mirror the Gaussian results, with a marginally significant relationship between phenology and demography overall, and negative but non-significant relationships in the group-specific analyses (Table 1; Fig. S1).

### Supplemental Visuals

**Table S1. Description of dataset.** Categories shown for overall meta-analytical dataset that includes 351 estimates from 89 studies (including all studies in all analyses data frames: Direct, Over Time & Experimental). Overall, 26 biomes were included in dataset, the top 10 summarized biomes are shown here.

| Category | # of Studies | # of Estimates | Estimate Percent |
|---|---|---|---|
| Study Type | | | |
| Experiments | 33 | 161 | 46.9% |
| Time Series | 43 | 142 | 41.4% |
| Comparisons of Historical Periods | 4 | 23 | 6.7% |
| Space for Time Substitutions | 2 | 13 | 3.8% |
| Extreme Year Comparisons | 1 | 4 | 1.2% |
| Experimental Treatments | | | |
| Experimental Warming | 15 | 70 | 53.0% |
| Snow Manipulations | 8 | 41 | 31.1% |
| Water Manipulation | 3 | 16 | 12.12% |
| Diet Manipulations | 2 | 4 | 3.0% |
| Phenology Manipulations | 1 | 1 | .06% |
| Study Organism | | | |
| Plants | 31 | 163 | 47.5% |
| Birds | 33 | 117 | 34.1% |
| Fish | 3 | 15 | 4.4% |
| Insects | 3 | 14 | 4.1% |
| Mammals | 7 | 12 | 3.5% |
| Reptiles | 3 | 12 | 3.5% |
| Amphibians | 3 | 10 | 2.9% |
| Hemisphere | | | |
| North Hemisphere | 78 | 327 | 95.3% |
| South Hemisphere | 5 | 16 | 4.7% |
| Environment | | | |
| Terrestrial | 73 | 304 | 88.6% |
| Aquatic | 5 | 21 | 6.1% |
| Terrestrial/Aquatic | 5 | 18 | 5.3% |
| Summarized Biome (Top 10) | | | |
| Mountain | 11 | 61 | 17.8% |
| Temperate Seasonal Forest | 19 | 59 | 17.2% |
| Woodland & Shrubland | 5 | 40 | 11.7% |
| Tundra | 10 | 35 | 10.2% |
| Grassland | 5 | 21 | 6.1% |
| Boreal Forest | 5 | 19 | 5.6% |
| Temperate Coniferous Forest | 2 | 19 | 5.6% |
| Wetland | 5 | 19 | 5.6% |
| Freshwater | 4 | 18 | 5.2% |
| Multiple Biomes | 5 | 18 | 5.2% |

**Table S2. Categories of Paper Exclusion**. A total of 175 studies were not able to be included in the meta-analyses. The top four categories are further described: **Unusable Data Format -** The paper only reported phenological metrics that are incompatible with our analyses, such as duration, mismatch metrics, and phenological progression system (number systems indicating different progressions of phenological stages). **Unable to Obtain Data -** The paper did not report necessary metrics, and the authors did not respond to contact attempts. **Environmental Focus -** The paper focused on how phenology and demography change due to environmental parameters instead of how phenology impacted demography or how they both changed over time. **Two Species focus -** The paper focused on how the phenology of one species affected the demography of a different species, but did not assess both parameters in one species.

| Reason Study Was Excluded | # of Studies | Percent |
|---|---|---|
| Unusable Data Format | 55 | 31.43% |
| Unable to Obtain Data | 33 | 18.86% |
| Environmental Focus | 26 | 14.86% |
| Two Species Focus | 25 | 14.29% |
| Unusable Analysis Type | 10 | 5.71% |
| Not Both Phenology & Demography | 8 | 4.57% |
| Unusable Variables | 7 | 4.00% |
| Other Focus | 3 | 1.71% |
| Replication of Dataset | 2 | 1.14% |
| Metrics Grouped for Multiple Species | 2 | 1.14% |
| Theory Paper (No Data) | 2 | 1.14% |
| Demography as Predictor | 1 | .006% |
| Lack of Control | 1 | .006% |

**Table S3. Analyses of Extent of Change in Phenology And Demography Across Time and Treatments.** All results came from rma.mv models from the metafor package with study ID as a random effect. Sample sizes (both the number of studies and the number of effect sizes included to account for studies with multiple effect sizes), the meta- estimate and 95% *CI* (calculated with robust error estimation) are shown. Analyses that are significant (p-value < .05) are bolded. $\tau^2$ represents the between-study variation in effect sizes, a 0 $\tau^2$ value indicates that no true variation in effect sizes has been detected and all variation is the result of estimation error.

| Data Source | Category | # of Studies | # Of Effects | Meta-Estimate | Meta-95% CI | | $\tau^2$ | df | p-value |
|---|---|---|---|---|---|---|---|---|---|
| Over Time | ***Phenology*** | ***13*** | ***45*** | ***-0.1245*** | ***-0.260*** | ***0.011*** | ***0.042*** | ***11.28*** | ***0.069*** |
| | Demography | 13 | 45 | 0.0005 | -0.004 | 0.005 | 0.001 | 8.37 | 0.826 |
| Experimental | **Phenology** | **26** | **132** | **-0.0435** | **-0.071** | **-0.016** | **0.005** | **24.87** | **0.003** |
| | Demography | 26 | 132 | -0.0801 | -0.187 | 0.026 | 0.062 | 24.05 | 0.134 |

**Table S4. Description of results of LMM's to understand the effect of phenological shifts on demographic vital rates for significant phenological shifts (95% CI does not cross 0).** All results came from LMM models constructed in lme4 with study ID as a random effect. Sample sizes (both the number of studies and the number of effect sizes included to account for studies with multiple effect sizes). Presented analyses all include more than five studies and 10 effect sizes (if groups were lower than these sample sizes, these were excluded from separated analyses but are included in the All category). Analyses that show marginally significant (p-value < .1) or significant (p-value < .05) are bolded. Treatment-Other was excluded from this analyses since it did not include enough studies. Historical comparisons and comparisons of extreme years are excluded from all experimental analyses.

| Data Frame | Category | # of Studies | # Of Effects | Estimate | 95% CI | | *t* | p-value |
|---|---|---|---|---|---|---|---|---|
| Over Time | All | 10 | 24 | -0.017 | -0.138 | 0.105 | -0.264 | 0.792 |
| | Animal-Birds | 7 | 19 | 0.032 | -0.105 | 0.152 | 0.531 | 0.595 |
| | Reproduction | 8 | 15 | -0.013 | -0.310 | 0.279 | -0.086 | 0.931 |
| Experimental | All | 22 | 61 | -0.050 | -1.421 | 1.344 | -0.072 | 0.942 |
| | Plant | 19 | 57 | -0.061 | -1.543 | 1.462 | -0.081 | 0.935 |
| | Reproduction | 18 | 50 | -0.235 | -1.748 | 1.304 | -0.304 | 0.761 |
| | **Survival & Size** | **6** | **11** | **3.545** | **-0.001** | **8.107** | **2.291** | **0.022** |
| | Treatment-Warming | 14 | 42 | -0.293 | -1.809 | 1.265 | -0.378 | 0.705 |
| | Treatment-Snow | 7 | 16 | 1.534 | -1.857 | 3.926 | 0.882 | 0.378 |

**Table S5.** Description of results of edgers regression test for regression test for funnel plot asymmetry. All analyses conducted are visualized in one table. Analyses that show marginally significant (p-value < .1) are italicized while analyses that show significant (p-value < .05) are bolded.

| Data Frame | Category | # of Studies | # Of Effects | Estimate | Sum. 95% CI | | z | p-value |
|---|---|---|---|---|---|---|---|---|
| Var- Gaussian | **All** | **26** | **80** | **0.0361** | **0.008** | **0.065** | **-4.200** | **0.0001** |
| | **Reproduction** | **22** | **66** | **0.0199** | **-0.012** | **0.052** | **-2.796** | **0.0052** |
| | **Survival & Size** | **5** | **14** | **0.6828** | **0.269** | **1.096** | **-5.687** | **0.0001** |
| | All Plant | 6 | 25 | -0.1172 | -0.801 | 0.567 | -0.787 | 0.4309 |
| | *All Animal* | *20* | *55* | *0.0187* | *-0.014* | *0.051* | *-1.696* | *0.0898* |
| | *Animal-Birds* | *15* | *37* | *0.0469* | *-0.041* | *0.135* | *-1.896* | *0.0579* |
| | Time Series | 19 | 60 | 0.0104 | -0.020 | 0.041 | -1.185 | 0.2361 |
| | **Experiments** | **6** | **17** | **0.4675** | **-0.252** | **1.187** | **-3.222** | **0.0013** |
| Var- Binomial | All | 16 | 29 | -0.0220 | -1.007 | 0.963 | 0.571 | 0.5678 |
| Var- Poisson | All | 13 | 28 | -0.0027 | -0.024 | 0.029 | -0.879 | 0.3796 |
| Time Series | *Phenology* | *13* | *45* | *0.0080* | *-0.036* | *0.052* | *-1.712* | *0.0868* |
| | Demography | 13 | 45 | 0.0001 | -0.010 | 0.011 | 0.026 | 0.9795 |
| Experimental | Phenology | 26 | 132 | -0.0207 | -0.062 | 0.020 | -0.569 | 0.5691 |
| | **Demography** | **26** | **132** | **0.1154** | **0.011** | **0.220** | **-2.767** | **0.0057** |

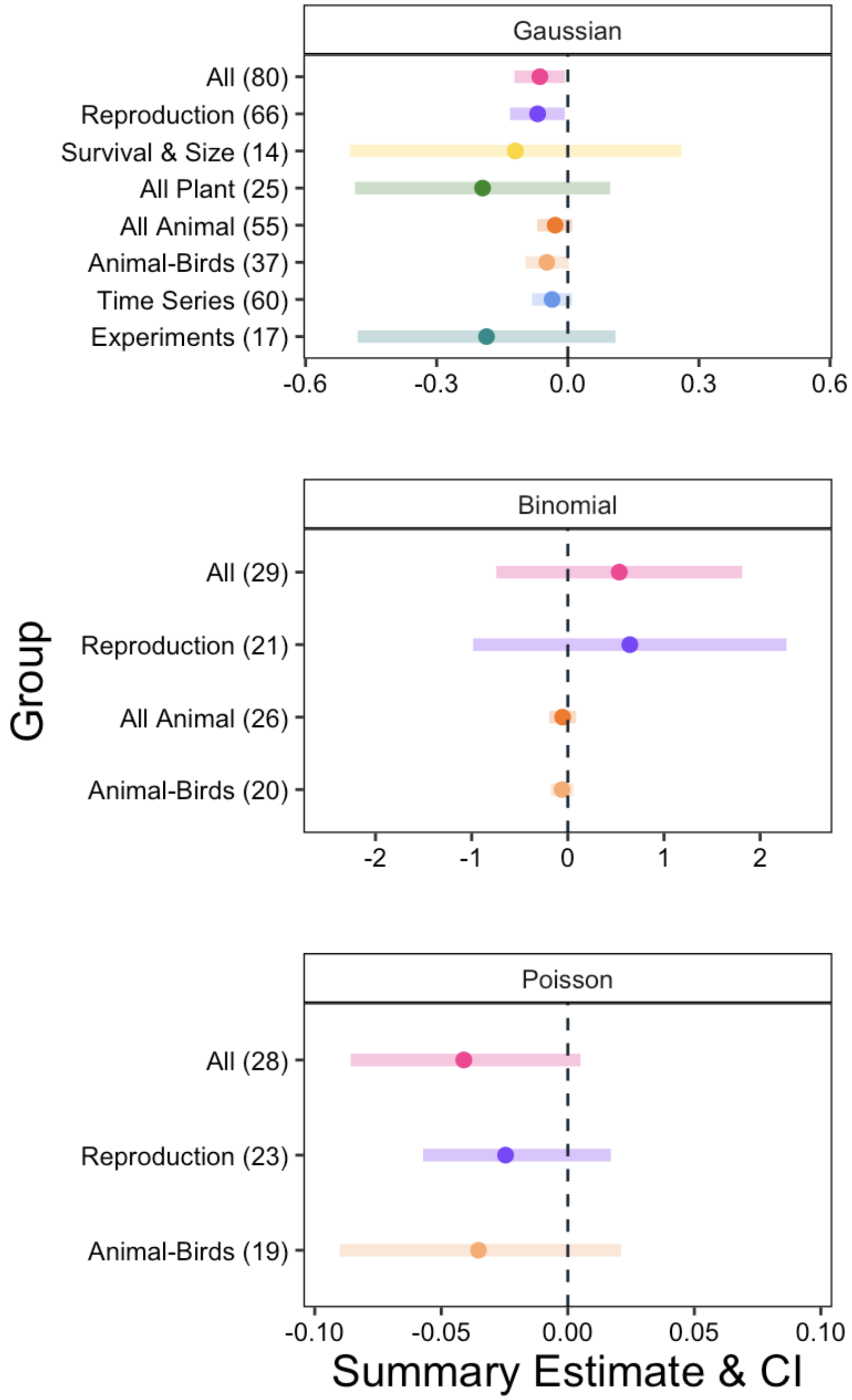


**Figure S1. Raw Estimates and SD for all direct studies split into separated error families colored by organism.** All citation information is shown on the y axis (including citation (first author, year, estimate number). Different x-axis scales are used for the different error family visuals.

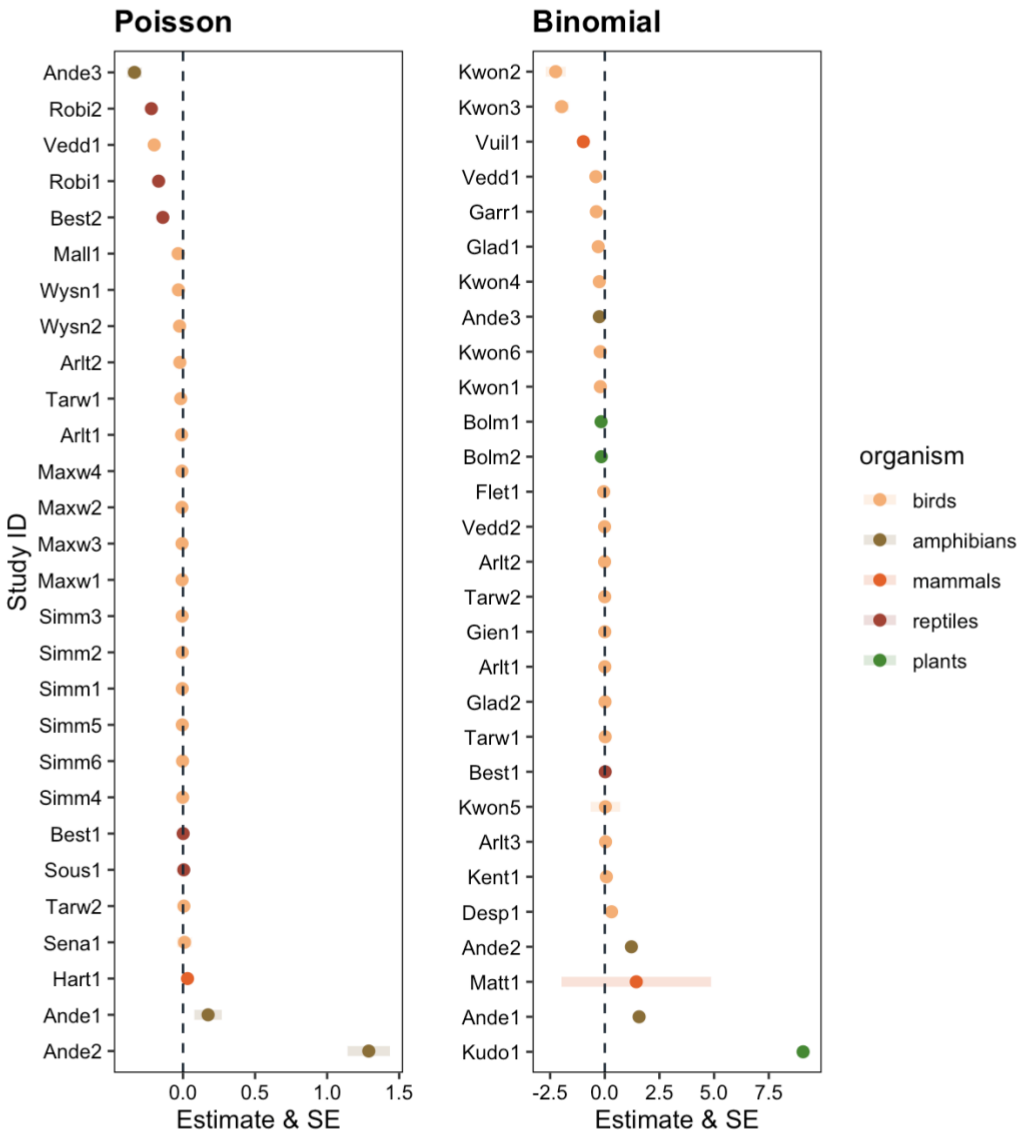


**Figure S2. Estimates and SE for all direct studies calculated using binomial and Poisson error families colored by organism.** Study ID is shown on the y axis (including first four letters of first author name and estimate number within the study. All estimates are visualized with error bars indicating SE although some are not visible due to large scale differences between studies.
